# PYK2 in the dorsal striatum of Huntington’s disease R6/2 mouse model

**DOI:** 10.1101/2024.01.18.576195

**Authors:** Omar Al-Massadi, Benoit de Pins, Sophie Longueville, Albert Giralt, Theano Irinopoulou, Mythili Savariradjane, Enejda Subashi, Silvia Ginés, Jocelyne Caboche, Sandrine Betuing, Jean-Antoine Girault

**Author notes:** **Corresponding authors:** Omar Al-Massadi and Jean-Antoine Girault (to whom correspondence should be addressed at the time of submission). Omar Al Massadi: Translacional Endocrinology group, Endocrinology section, Instituto de Investigación Sanitaria de Santiago de Compostela, Complexo Hospitalario Universitario de Santiago (IDIS/CHUS), Santiago de Compostela. CIBER Fisiopatología de la Obesidad y Nutrición (CIBERobn), Santiago de Compostela, Spain. Benoit de Pins, Department of Plant and Environmental Sciences, Weizmann Institute of Science, Rehovot, Israel.

## Abstract

Huntington’s disease (HD) is a devastating disease due to autosomal dominant mutation in the *HTT* gene. Its pathophysiology involves multiple molecular alterations including transcriptional defects. We previously showed that in HD patients and mouse model, the protein levels of the non-receptor tyrosine kinase PYK2 were decreased in the hippocampus and that viral expression of PYK2 improved the hippocampal phenotype. Here, we investigated the possible contribution of PYK2 in the striatum, a major brain region altered in HD. PYK2 mRNA levels were decreased in the striatum and hippocampus of R6/2 mice, a severe HD model. PYK2 protein levels were also decreased in the dorsal striatum of R6/2 mice and in the putamen of human patients. PYK2 knockout by itself did not result in motor symptoms observed in HD mouse models. Yet, we examined whether PYK2 deficiency participated in the R6/2 mice phenotype by expressing PYK2 in the dorsal striatum using AAV vectors. With an AAV1/*Camk2a* promoter, we did not observe significant improvement of body weight, clasping, motor activity and coordination (rotarod) alterations observed in R6/2 mice. With an AAV9/*SYN1* promoter we found an improvement of body weight loss and a tendency to better rotarod performance. DARPP-32 immunofluorescence was increased following AAV9/*SYN1*-PYK2 injection compared to AAV9/*SYN1*-GFP, suggesting a possible partial beneficial effect on striatal projection neurons. We conclude that PYK2 mRNA and protein levels are decreased in the striatum as in hippocampus of HD patients and mouse models. However, in contrast to hippocampus, striatal viral expression of PYK2 has only a slight effect on the R6/2 model striatal motor phenotype.

**Highlights:**

- Huntington’s disease is a lethal genetic disease altering striatum, cortex, and hippocampus
- Restoring PYK2 levels in hippocampus improved hippocampal phenotype of a Huntington mouse model
- We show that PYK2 levels are decreased in the striatum of R6/2 mice and human patients
- Viral expression of PYK2 in the striatum has only a small effect on R6/2 mouse model motor phenotype but improves weight loss

## 1. Introduction

Huntington’s disease (HD) is an autosomal dominant inherited disease caused by trinucleotide repeat expansion in the Huntingtin gene (*HTT*) (Collaborative, 1993). The prevalence of HD is 3-12 cases per 100,000 people of European ancestry (Evans et al., 2013; Pringsheim et al., 2012). HD patients display a broad symptomatology affecting cognitive function and behavior. These patients also develop major motor impairments including gait disturbances, lack of coordination, choreic and jerky body movements. Another hallmark of the disease is the loss of body weight despite efforts to maintain high calorie diet (Marder et al., 2009). Although great progress has been achieved in understanding the molecular and cellular mechanisms in HD (Saudou and Humbert, 2016), virtually no therapeutic options exist to treat this disease. Thus, the identification of new molecular targets that could help in the fight against HD is urgently desired.

Since the discovery of the *HTT* gene mutation numerous genetic mouse models were genetically engineered (Figiel et al., 2012; Heng et al., 2008; Pouladi et al., 2013). Many motor and cognitive deficits endured by HD patients are recapitulated in these preclinical mouse models, which are useful tools to uncover pathophysiology and to develop potential therapeutic strategies (Crook and Housman, 2011; Gil and Rego, 2009; Switonski et al., 2012). Among the genetic mouse lines commonly used to study HD pathogenesis, the R6/1 and R6/2 mice are transgenic for the first exon of the human HTT gene with amplified CAG repeats (Brooks and Dunnett, 2015; Farshim and Bates, 2018; Stricker-Shaver et al., 2018). The R6/2 line displays an accelerated phenotype and develops neurological abnormalities, including loss of coordination, abnormal gait, hind limb clasping behavior neuropathology, loss of body weight and premature death (Brooks and Dunnett, 2015; Farshim and Bates, 2018; Stricker-Shaver et al., 2018). R6/2 mice develop symptoms at the age of 5 weeks and can live until 14 weeks whereas the R6/1 mice model has a longer life expectancy and less severe symptomatology (Brooks and Dunnett, 2015; Farshim and Bates, 2018; Stricker-Shaver et al., 2018).

The most prominent neuronal change in HD is a pathological deterioration in the dorsal striatum (DS) (Klapstein et al., 2001; Levine et al., 1999; Vonsattel and DiFiglia, 1998). The DS receives information from all areas of the cerebral cortex and its role in motor behavior is altered in HD (Estrada-Sanchez and Rebec, 2013). Striatal projection neurons (SPNs) are medium-size spiny GABAergic neurons, which account for more than 90% of all striatum neurons and are particularly vulnerable in HD, undergoing progressive degeneration and massive cell loss over the course of the disease (Klapstein et al., 2001; Levine et al., 1999; Vonsattel and DiFiglia, 1998). Therefore, the DS appears as a key target area in the disease.

PYK2 is a non-receptor tyrosine kinase that can be activated by Ca^+2^ and is highly expressed in the brain, specifically in forebrain principal neurons [(Menegon et al., 1999) and recent review in (de Pins et al., 2021)]. Previous studies determined that PYK2 is involved in synaptic plasticity (Bartos et al., 2010; Giralt et al., 2017; Huang et al., 2001; Mastrolia et al., 2021) and its gene, *PTK2B*, is a risk locus for Alzheimer’s disease (Lambert et al., 2013). Our group previously showed that i) PYK2 protein levels are decreased in the hippocampus of HD patients and R6/1 mice; ii) *Ptk2b* knockout (PYK2-KO) mice hippocampal phenotype resembles that of R6/1 mice; iii) PYK2 viral expression in the hippocampus of R6/1 mice improves memory deficits and some synaptic markers (Giralt et al., 2017).

Given the key role of the striatum in HD and our previous data suggesting a contribution of PYK2 deficit in the hippocampus in HD phenotype, we examined the possible alteration of PYK2 in the DS. We measured PYK2 mRNA and protein in R6/2 mice and patients, we studied the motor function of PYK2-KO mice and we investigated the effects of AAV-mediated PYK2 expression in the DS of R6/2 mice.

## 2. Material and Methods

### 2.1. Mice

Wild type mice were male C57BL/6JRj mice provided by Janvier Laboratories (France). R6/2 [B6CBA-Tg (HDexon1) 62Gpb/1J] mice, which express exon 1 of the human mutant Huntington’s disease gene containing 150 CAG repeats, under the control of the human HTT (IT15) promoter, were obtained from Jackson Laboratories by crossing ovarian transplant hemizygous females with B6CBAFI/J males. Mice used in this study were males and females from the first offspring and the genotype was verified by polymerase chain reaction (PCR) using genomic DNA extracted from the tail. The number of CAG repeat length varies very little in the progeny from the first generation (http://chdifoundation.org/wp-content/uploads/HD_ Field_Guide_040414.pdf), and can therefore be considered around 150 CAG for all the mice that were used in the study. PYK2-KO male mice, which bear a deletion of exons 15 to 18 in the *Ptk2B* gene which codes PYK2, were produced by our laboratory as previously described (Giralt et al., 2017; Giralt et al., 2016). WT littermates and mutant mice from either sex were fed *ad libitum* and housed in groups under conditions of controlled temperature (23°C) and illumination (12-hour light/12-hour dark cycle). Animal experiments and handling was in accordance with ethical guidelines of Declaration of Helsinki and NIH, (1985-revised publication no. 85-23, European Community Guidelines), and French Agriculture and Forestry Ministry guidelines for handling animals (decree 87849, license A 75-05-22) and approval of the Charles Darwin ethical committee (APAFIS#8861-2016111620082809).

### 2.2 Viral constructs and stereotaxic injection

For expression of PYK2 in the DS, 4-week-old R6/2 and WT mice were stereotaxically injected with AAV vectors. Groups of animals received AAV1.Camk2a-eGFP-2A-mPTK2B (referred to below as AAV-Camk2a-PYK2) expressing PYK2 and GFP separated by a T2A cleavable link (Vector Biolabs Malvern, PA, USA). As control we used AAV9.Camk2a-eGFP.WPRE.rBG (AAV-Camk2a-GFP) expressing only GFP. In different groups of mice we used AAV9-hSYN1-eGFP-2A-mPTKB2B (AAV-SYN1-PYK2) and as control AAV9-hSYN1-eGFP-WPRE (AAV-SYN1-GFP) from University of Pennsylvania Perelman School of Medicine AAV core facility. Mice were anesthetized by an intraperitoneal injection of ketamine, 15 mg/kg and xylazine, 3 mg/kg body weight, and placed in a stereotaxic frame (Kopf Instruments). The DS was targeted bilaterally as previously reported (Boussicault et al., 2016), using a 32-gauge needle connected to a 1-ml syringe (Neuro-Syringe, Hamilton) at stereotaxic coordinates 0.5 mm rostral to the bregma, 2.1 mm lateral to midline, and 3.35 mm ventral to the skull surface. The rate of injection was 0.15 μl/min for 13 min with a total volume of 2 µl injected per striatum. After AAV infusion the needle was kept in place for an additional 5 minutes. After the procedure, the incision was closed with black braided non absorbable silk suture (Ethicon, Puerto Rico USA), and mice were placed 2 h for careful monitoring in a heated cage until they recovered from anesthesia then returned to their home cage for 2 weeks before starting analyses of behavior. Analgesia was used during the two days following surgery by subcutaneous injections of a non-steroidal anti-inflammatory drug (flunixin, 4 mg kg^−1^, Cronyxin, Bimeda).

### 2.3 Behavioral phenotyping

#### Open Field

The open field apparatus was a dimly lit (60 lx) white square arena (40 × 40 × 40 cm in length, width, and height respectively). Animals were placed at the center and allowed to explore freely for 15 min. Spontaneous locomotor activity was recorded and analyzed with a video tracker (Viewpoint, Newcastle Upon Tyne, UK), a software that automatically detects rodent movements and behaviors (Giralt et al., 2017).

#### Circular maze

Locomotor activity was measured in a circular corridor with four infrared photoelectric beams placed at 90° angles (Imetronic, Pessac, France) in a low luminosity environment. Mice were placed in the maze for 90 min and the spontaneous locomotion measured in ¼ turns (consecutive interruptions of two adjacent beams) and the number of rearings in 5-min time bins.

#### Balance beam

The balance beam test is a sensorimotor integration test evaluating balance and motor skills. The test measures the time taken by the mouse to reach a safe enclosed platform, walking over a narrow and elevated beam. A bright light illuminates the start area and a close 20 x 20 cm square escape box is located at one end of the beam. During the two consecutive days of testing, each mouse was given two trials with a minimum of 10 min between the trials. One day before the tests, a habituation session was given to familiarize the mouse with the beam. For each trial, the mouse was placed in the center of the beam facing one of the platforms and then released. Latency to traverse the beam was recorded.

#### Vertical Pole Test

The vertical pole test measures the balance, motor coordination, and vertical orientation capability of mice. The mouse was placed head downward in the middle of a round wooden pole (50-cm height, 1-cm diameter). The pole was wrapped with cloth tape for improved traction and preventing sliding. The pole was held in a horizontal position, and was then gradually shifted to a vertical position. The time until the mouse stayed or walked up or down the pole was measured with a stopwatch. The cut off time of the test was 60 seconds to avoid exhaustion. The time of latency to fall was calculated for each mouse. All mice were habituated to the task one day before the test, followed by a one-day test. During testing, the following scores were applied: Falling the pole immediately or when trying to turn or descend at any time (scored 0); 0-20 seconds of latency (scored 1); 21-40 seconds of latency (scored 2); 41-60 seconds of latency (scored 3); More than 60 seconds of latency (scored 4); Sliding down the pole immediately (scored 5); Freezing and never descending the pole (scored 6).

#### Rotarod

Rotarod was used to evaluate the motor coordination (Boussicault et al., 2016). Mice were tested over three consecutive days, each daily session included a 5 min training trial at 4 rpm on the rotarod apparatus (LE8200, Bioseb). One hour later, the animals were tested for three consecutive accelerating trials of 5 min with the rotarod speed linearly increasing from 4 to 45 rpm over 300 s. The latency to fall from the rod and the speed at fall were recorded for each trial. Mice remaining on the rod for more than 300 s were removed and their time scored as 300 s. Data from the training trial were not included.

#### Hind limb clasping

Mice were tested at 6, 9, and 12 weeks of age. They were suspended by the tail for 30 s and the clasping phenotype was graded according to the following scale: level 0, no clasping; level 1, clasping of the forelimbs only or both fore-and hind limbs once or twice; and level 2, clasping of both fore and hind limbs more than three times or more than 5 s (Boussicault et al., 2016).

### 2.4 Immunohistofluorescence

Mice received a lethal dose of pentobarbital (500 mg/kg, i.p., Dolethal, Vetoquinol) and perfused transcardially with 40 g/L paraformaldehyde in 0.1 M sodium phosphate buffer pH 7.4. Brains were post-fixed overnight at 4°C and cut into 30 μm-thick free-floating coronal sections with a vibrating microtome (Leica, VT1000). All the following steps were done with gentle shaking. Sections were washed three times in phosphate-buffered saline (PBS), permeabilized in PBS-T (PBS + 3 µl/ml Triton X-100) and 30 µl/ml normal goat serum (Pierce Biotechnology, Rockford, IL, USA) for 60 min at room temperature, and washed three times. Brain slices were incubated overnight at 4°C in the presence of primary antibodies in PBS-T: rabbit PYK2 antibody (1/500, Sigma #07M4755, Chemical Co.,St Louis, MO, USA), chicken GFP antibody (Abcam, EPR14104, ab300643), and DARPP-32 (1/5 000; gift from Paul Greengard, The Rockefeller University). Sections were then washed three times and incubated for 2 h at room temperature with fluorescent secondary antibodies: Cy3 goat anti-rabbit (1/200) and/or AlexaFluor 488 goat anti-mouse (1/200; both from Jackson ImmunoResearch, West Grove, PA, USA). No signal was detected in control sections incubated in the absence of the primary antibody. Images of the whole slices (10x) were acquired on a Leica DM6000 microscope and quantified with Imaris software at the *Institut du Fer à Moulin Cell and tissue imaging* facility.

### 2.5. RT-qPCR

RNA was extracted using Trizol^®^ reagent (Invitrogen) according to the manufacturer’s instructions. The RNA was quantified with a Nanodrop 1000 spectrophotometer. Five hundred ng of total RNA were used for each reaction. Reverse transcription, cDNA synthesis was performed using the SuperScript™ First-Strand synthesis System (Invitrogen) and random primers. Negative control reactions, containing all reagents except the sample, were used to ensure specificity of the PCR amplification. Real-time polymerase chain reaction was carried out using a fluorescent temperature cycler (TaqMan®; Applied Biosystems, Foster City, CA, USA) following the manufacturer’s instructions. The oligonucleotide specific primers are shown in Supplemental Table 1. Results were quantified and normalized to a house-keeping gene with delta-delta-CT (ddCT) method and expressed in comparison with the average value for the control group. All samples were run in duplicate and the average values calculated.

### 2.6. Immunoblot analysis

*Mouse brain samples.* Immunoblotting was performed as described (Giralt et al., 2017; Giralt et al., 2018; Montalban et al., 2019). Briefly, animals were euthanized by CO_2_ inhalation and the DS rapidly dissected on ice. Protein lysates from DS (10 μg) were subjected to sodium dodecyl sulfate–polyacrylamide gel electrophoresis, electrotransferred onto nitrocellulose membranes (GE Healthcare, LC, UK), and probed with rabbit PYK2 antibodies (diluted 1/1 000, 074M4755, Sigma). After several washes in TBS-Tween, blots were incubated with secondary anti-rabbit IgG IRdye800CW antibodies (1:2,000, Rockland Immunochemicals, USA). Secondary antibody binding was detected by Odyssey infrared imaging apparatus (Li-Cor Inc., Lincoln, NE). Membranes were also incubated with an antibody for β-actin or α-tubulin, either simultaneously or successively, which was used as a loading control to normalize PYK2 levels. Molecular weight were checked using a Prestained Protein Ladder (#06P-0111, Euromedex).

#### Post-mortem human brain

Postmortem putamen tissue samples from control and HD patients were obtained from the Neurological Tissue Bank (Biobanc-HC-IDIBAPS), Barcelona, Spain with the approval of the local ethics committee at recruitment and an informed written consent (Martin-Flores et al., 2020). They included six controls (two females, four males aged 81.2 ± 5.0 years, mean ± SD, post-mortem interval, 11.9 ± 8.8 h) and six patients with HD grades 1-3 (all males, age 68.2 ± 12.3 years, post-mortem intervals, 8.8 ± 4.2 h, see **Table 1**). Immunoblotting was done as described for mouse samples.

**Table 1:**
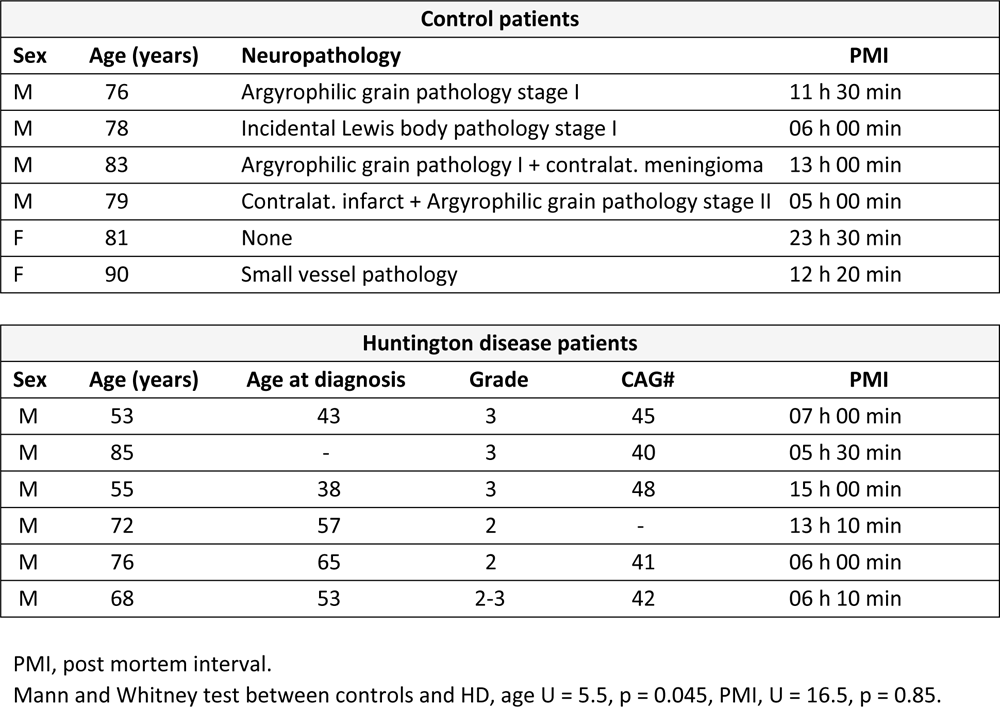
Human post-mortem samples.

### 2.7 Statistical analyses

Analyses were done using GraphPad Prism 6. After assessing normality of data distribution with d’Agostino-Pearson and Shapiro-Wilk tests, parametric analysis was conducted. Two-group comparisons were done with two-tailed unpaired Student’s t test and 3 groups analyzed using one-or two-way ANOVA, followed by Holm-Sidak post hoc test for multiple comparisons. If the distribution was significantly different from normal or if the sample size of one or more data sets was too small (<7), non-parametric analysis was applied. Non-parametric comparisons of 2 groups were done with Mann-Whitney test and 3 groups or more with Kruskal-Wallis test followed by Dunn or Mann-Whitney test as indicated. Significance threshold was 0.05. Detailed statistical results are presented in **Supplementary Table 1**.

## 3. Results

### 3.1. PYK2 mRNA levels are decreased in the hippocampus and striatum but not in the cerebral cortex of R6/2 mice

The expression of PYK2 mRNA in the brain was previously investigated by *in situ* hybridization (Menegon et al., 1999). To obtain more quantitative results, we performed RT-PCR using samples from hippocampus, striatum, and cerebral cortex of 3-4 month-old WT male mice. We confirmed the data of Menegon et al., who found that the hippocampus is the brain region with the highest PYK2 mRNA expression (**Figure 1A**). PYK2 mRNA levels in the striatum were intermediate between those in the cerebral cortex and in the hippocampus (**Figure 1A**, **Supplementary Table 1**). These measurements indicated that PYK2 may play a role in the striatum in the context of HD. We then investigated whether PYK2 mRNA levels were altered in the R6/2 HD mouse model and measured PYK2 mRNA using RT-qPCR in the cerebral cortex, hippocampus, and striatum. In the cortex of 12-week-old R6/2 mice, because PYK2 mRNA levels did not differ significantly from those in WT controls (**Figure 1B**, **Supplementary Table 1**), we did not investigate younger ages. In the hippocampus and striatum, we measured PYK2 mRNA levels at 4, 6 and 12 weeks of age (**Figures 1C and D**). In both regions PYK2 mRNA levels were decreased in R6/2 mice at 6 and 12 weeks as compared to WT mice, but not at 4 weeks (**Figure 1C and D**, **Supplementary Table 1**). These data showed an earlier loss of PYK2 transcripts in the striatum and hippocampus of R6/2 mice than in the cortex although the three brain regions display early metabolic alterations (Mochel et al., 2012). They indicated that the previously reported deficit in PYK2 protein in the hippocampus of an HD mouse model (Giralt et al., 2017) is likely to result from a decrease in mRNA. In addition, they suggested a possible contribution of PYK2 in the phenotype of R6/2 mice related to striatal function.

**Figure 1:**
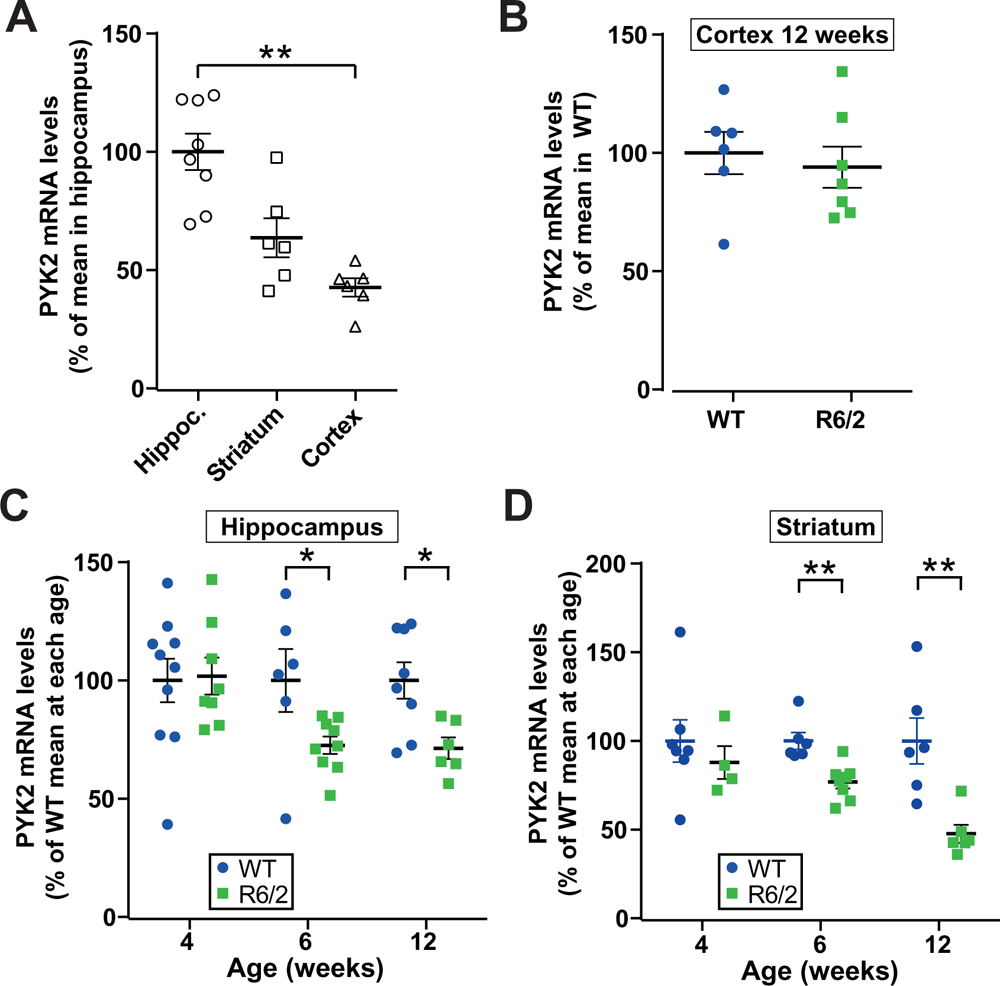
PYK2 mRNA levels are decreased in hippocampus and striatum of R6/2 mice. PYK2 mRNA levels were quantified by RT-qPCR. **A**) PYK2 mRNA in the hippocampus, striatum and cerebral cortex of 12-week old WT male C57Bl/6 mice (6–8 animals per group). Data are expressed as % of the mean in hippocampus, Kruskal-Wallis test, p < 0.0001, Dunn’s multiple comparisons test. **B**) PYK2 mRNA levels in the cerebral cortex of 12-week WT and R6/2 mice (6-7 animals per group). Mann Whitney test, not significant (ns). **C**) PYK2 mRNA levels in the hippocampus of WT and R6/2 mice at the indicated ages (6-10 animals per group). Kruskal-Wallis test, p = 0.0085, Mann-Whitney test, 4 w, ns, 6 w, p = 0.036, 12 w, p = 0.02. **D**) PYK2 mRNA levels in the DS of WT and R6/2 mice at the indicated ages (4-8 animals per group). Kruskal-Wallis test, p = 0.0017, Mann-Whitney test, 4 w, ns, 6 w, p = 0.0047, 12 w, p = 0.0043. (**A-D**) Means ± SEM are indicated, * p < 0.05, ** p < 0.01, detailed statistical results in **Supplementary Table 1**.

### 3.2. Striatal PYK2 protein levels are decreased in the striatum of R6/2 mice and HD patients

Since changes in mRNA levels can be inconsistent with the levels of the corresponding protein, we investigated PYK2 protein levels in the striatum of R6/2 mice using immunoblotting. PYK2 protein levels were not changed at 6 weeks of age but decreased at 9 and 12 weeks (**Figure 2A-B**, **Supplementary Table 1**). We then investigated PYK2 protein levels in post-mortem samples of the putamen of HD patients compared with controls (**Table 1**). PYK2 levels were reduced in HD patients (controls, 100 ± 9.7%, HD, 63.6 ± 11.8%, mean ± SEM, 6 subjects per group, **Figure 2C-D**, **Supplementary Table 1**). These results open the possibility of a potential role for a PYK2 deficit in the striatum of HD patients in the pathophysiology and symptomatology, as previously observed in the hippocampus (Giralt et al., 2017).

**Figure 2:**
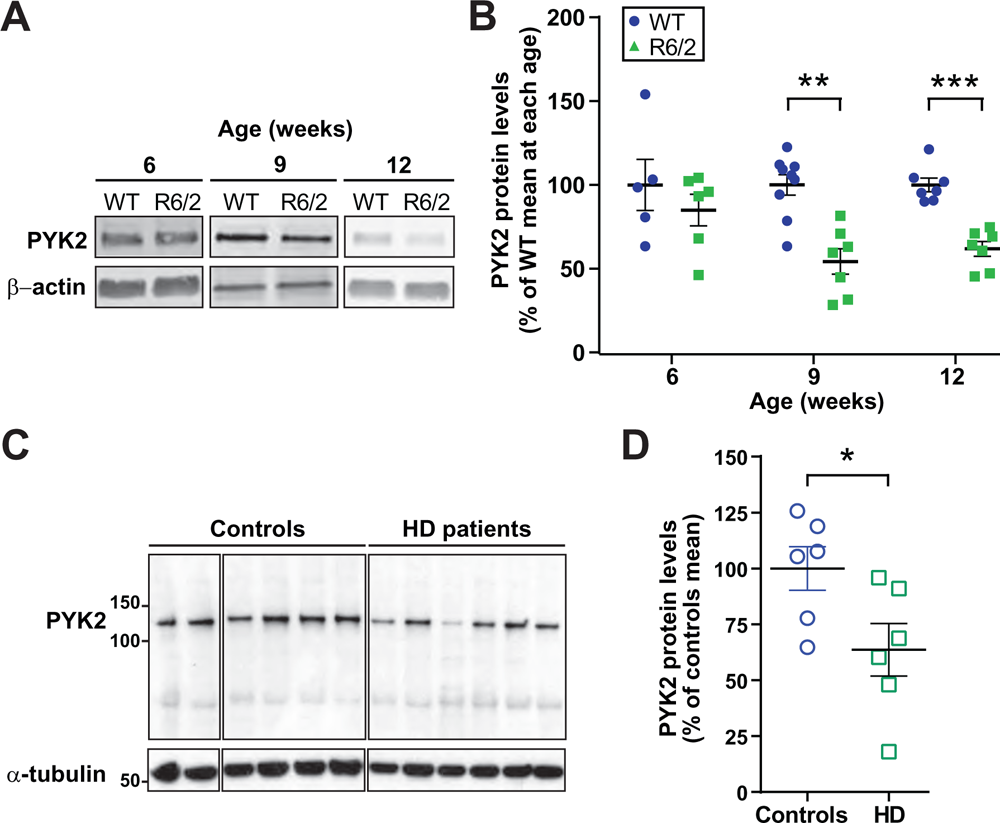
PYK2 protein levels are decreased in the striatum of R6/2 mice and HD patients. **A**) PYK2 protein levels were measured by immunoblotting in the striatum of WT and R6/2 mice at the indicated ages. β-actin was used as a loading control. **B**) Data as in (**A**) were quantified by densitometry (5-9 animals per group). PYK2 levels were normalized to the loading control and data expressed as % of the mean in WT at each age. Kruskal-Wallis test, p = 0.0007. Mann-Whitney test, 4 w, ns, 6 w, p = 0.0012, 12 w, p = 0.0006. **C**) PYK2 protein levels were measured by immunoblotting in the putamen of human subjects deceased with **HD** or with other conditions (**Table 1**). α-tubulin was used as a loading control. Data were quantified, normalized and expressed as % of the mean in controls (6 controls and 6 HD). Mann-Whitney test, p = 0.041. (**A-B**) Means ± SEM are indicated,* p < 0.05, ** p < 0.01, *** p < 0.001, detailed statistical results in **Supplementary Table 1**. Uncropped immunoblots for all samples are shown in **Supplementary** Figure 1.

### 3.3. PYK2-KO mice exhibit normal motor behavior

Because PYK2 mRNA and/or protein were decreased in the striatum of HD patients and R6/2 mouse model, we next examined whether the absence of PYK2 was able by itself to induce motor deficits. For this purpose, we used 3-4 month-old male PYK2-KO mice. We first measured their locomotor activity in an open field and found that PYK2-KO mice traveled the same distance as WT littermates (**Figure 3A**, **Supplementary Table 1**). We then compared the sensori-motor coordination of the PYK2-KO and WT littermate mice in the balance beam, with two trials for each mouse on two consecutive days (**Figure 3B**, **Supplementary Table 1**). We did not observe any difference between the two genotypes in this test. We also compared the PYK2-KO and WT littermate mice performance in the vertical pole test, which evaluates balance, motor coordination, and vertical orientation capabilities. PYK2-KO mice did not exhibit deficits in this test (**Figure 3C**, **Supplementary Table 1**). We also looked for a characteristic response exhibited by HD animal models, hind limb clasping (Boussicault et al., 2016; Mangiarini et al., 1996). We did not observe this response in PYK2-KO mice (**Figure 3D**). The absence of patent motor phenotype in PYK2-KO animals was consistent with our previous observation of normal performance of PYK2-KO mice on the accelerating rotarod (de Pins et al., 2020) and showed that the complete lack of this protein is not sufficient to visibly alter motor function in young mice. These results suggested that the partial decrease in PYK2 mRNA and protein levels observed in the striatum of HD mouse models or patients is unlikely to contribute by itself to the motor alterations they display. Yet, they do not rule out the possibility that the PYK2 deficit synergizes with other molecular defects in HD and plays an aggravating role.

**Figure 3:**
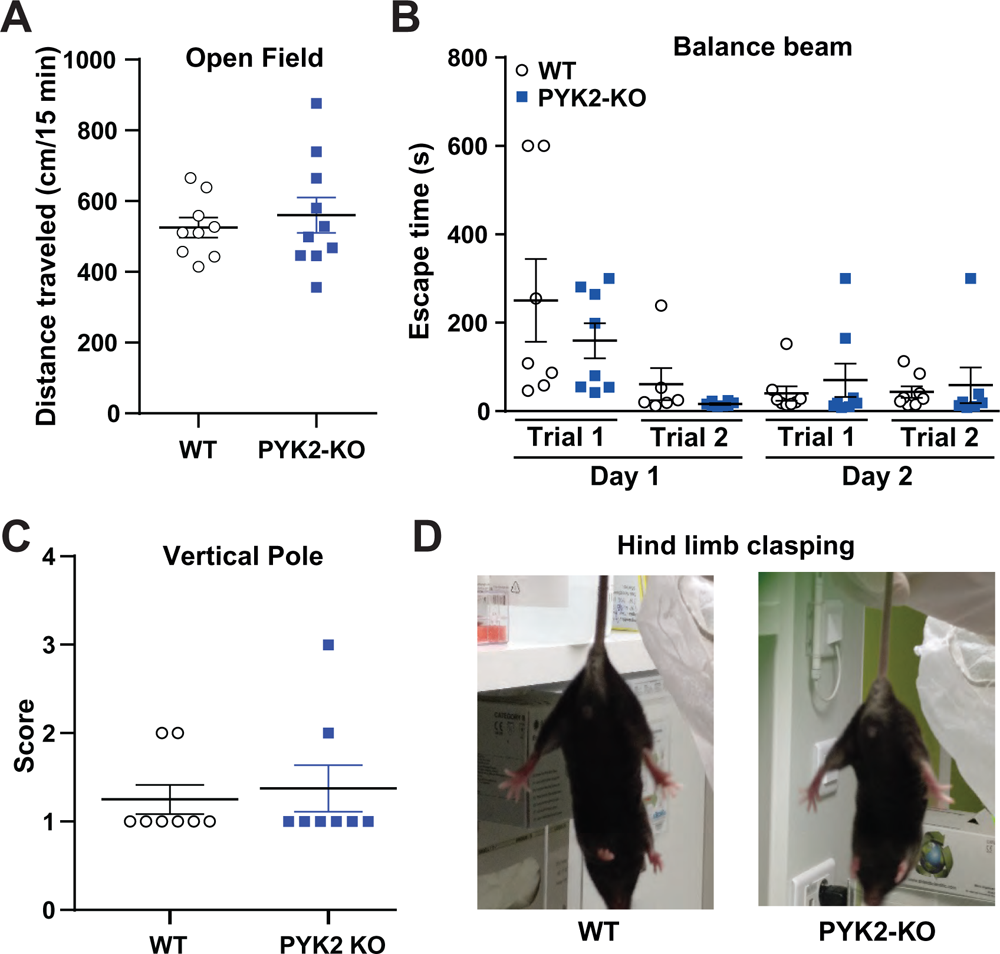
Motor behavior is not altered in PYK2-KO mice. **A**) The spontaneous locomotor activity of 3-6-month-old male WT and PYK2-KO littermate mice was studied by video tracking for 15 min in an open field. Unpaired 2-tailed Student’s t test, ns (9-10 mice per group). **B**) Motor skill was evaluated as the time to reach the escape zone on a balance beam during 2 trials on 2 consecutive days (6-8 mice per group). Kruskal-Wallis test, p = 0.0007, Dunn multiple comparisons test, no difference between genotypes. **C**) Motor skill was evaluated on a vertical pole by visually scoring (8 mice per group). Mann-Whitney test, ns. **D**) Examples of hind limb position when mice were held by the tail showing the absence of clasping in WT and PYK2-KO mice. (**A-C**) Means ± SEM are indicated. Detailed statistical results in **Supplementary Table 1**.

### 3.4. Lack of effect of AAV-mediated PYK2 striatal expression under *Camk2a* promoter in R6/2 mice

Since the expression levels of PYK2 in the striatum of different mouse models and in the putamen of HD patients were decreased (**Figure 2**) and because re-expression of PYK2 in the hippocampus had beneficial effects on the symptoms linked to this brain region (Giralt et al., 2017), we asked whether expressing PYK2 in the striatum of R6/2 mice, could partly improve their phenotype. We used AAVs expressing PYK2 and GFP or GFP alone under a mouse *Camk2a* promoter. We first tested the method in WT mice by injecting AAV-*Camk2a*-GFP in one DS and AAV-*Camk2a*-PYK2 in the contralateral striatum (**Fig. 4A**). Two weeks after the injection, GFP expression demonstrated the spreading of viral transduction (**Figure 4A**). Immunostaining in WT mice (**Figure 4A**) showed that AAV-*Camk2a*-PYK2 injection raised PYK2 levels by comparison to AAV-*Camk2a*-GFP.

**Figure 4:**
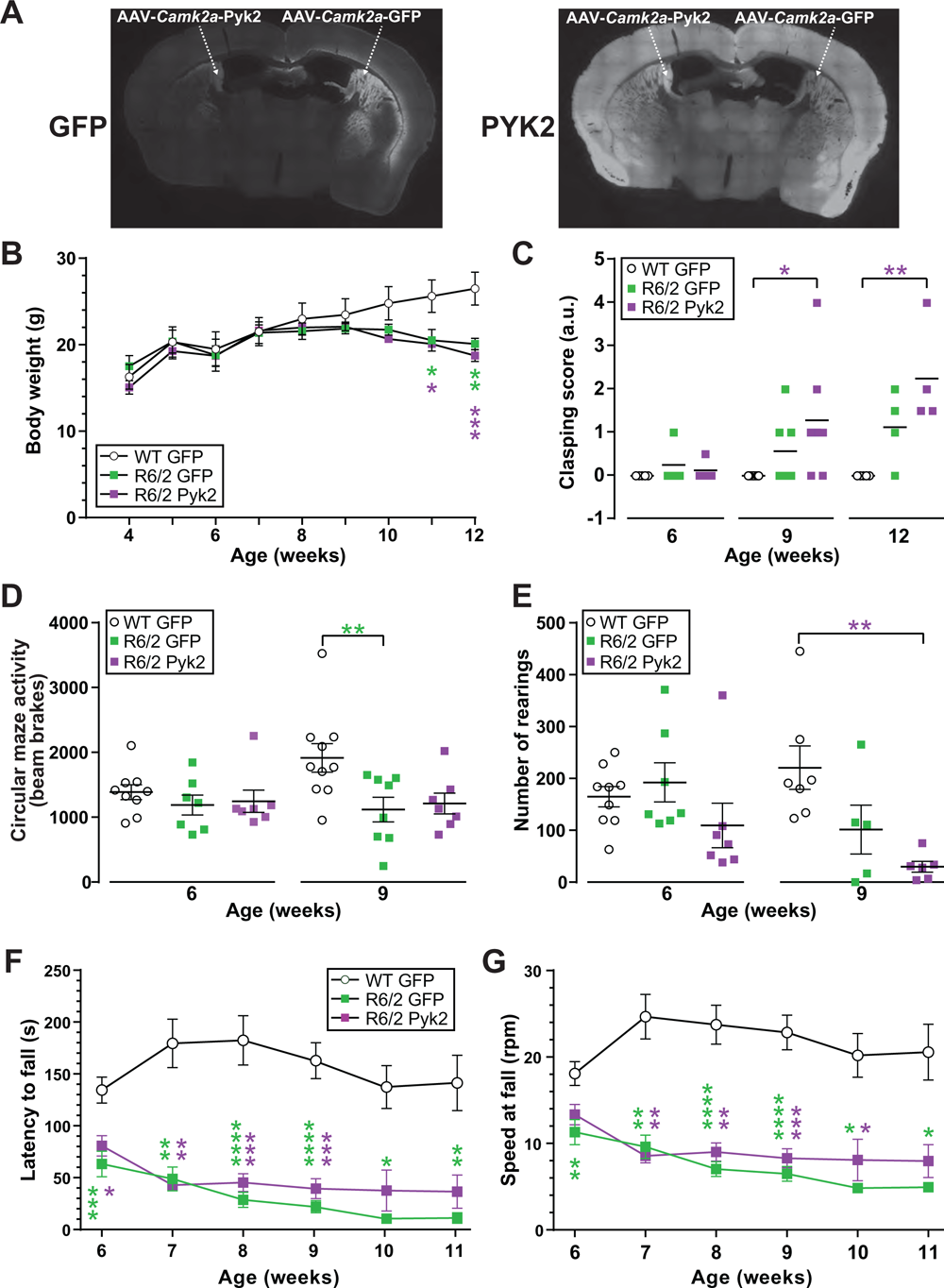
Effects of AAV-*Camk2a*-PYK2 transduction in the DS of R6/2 mice. **A**) AAV-*Camk2a*-PYK2, expressing PYK2 and GFP, was stereotactically injected in one striatum of a WT mouse and AAV-*Camk2a*-GFP, expressing GFP, in the contralateral striatum. Coronal brain section were immunostained for GFP (left picture) and PYK2 (right picture). **B**) R6/2 mice received a bilateral injection of either AAV-*Camk2a*-PYK2 or AAV-Camk2a-GFP in the DS and WT mice a bilateral injection of AAV-Camk2a-GFP. Their body weight was measured every week. Two way repeated measures ANOVA, interaction, F_(16,80)_ = 8.79, p < 10^-4^, age effect, F_(8,80)_ = 45.0, p < 10^-4^, mouse/AAV group effect, F_(2,10)_ = 1.0, p = 0.39 (4-5 mice per group). Asterisks indicate significant post hoc Holm-Sidak multiple comparisons test. **C**) Clasping responses were scored in the 3 groups at the indicated ages (4-7 mice per group and age). Kruskal-Wallis test, p = 0.0006. Horizontal bars are means. **D-E**) The motor behavior of the 3 groups of mice was studied in a circular maze at 6 and 9 weeks (7-10 mice per group and age). General locomotor activity (number of beam breaks, **D**) and the number of rearings (**E**) were measured. Kruskal-Wallis test, (**D**) 6 weeks, ns, 9 weeks p = 0.018, (**E**) 6 weeks, p = 0.038, 9 weeks p = 0.002, Dunn’s multiple comparisons test. **F-G**) The motor coordination was evaluated every week from 6 to 11 weeks (w) in a rotarod test by measuring the latency to fall (**F**) and the speed at fall (**G**) (4-14 mice per group and age). Kruskal-Wallis test at each age, (**F**) 6 w, p = 0.0007, 7 w, p = 0.0006, 8 w p < 10^-4^, 9 w p < 10^-4^, 10 w, p = 0.013, 11 w, p = 0.0091, (**G**) 6 w, p = 0.0047, 7 w, p < 10^-4^, 8 w p < 10^-4^, 9 w p < 10^-4^, 10 w, p = 0.0028, 11 w, p = 0.0011. (**B**, **D-G**) Means ± SEM are indicated. (**C-G**) Post hoc Dunn’s multiple comparisons test. Significant differences are indicated by asterisks, green, R6/2 AAV-*SYN1*-GFP vs WT AAV-*SYN1*-GFP, violet, R6/2 AAV-*SYN1*-PYK2 vs WT AAV-*SYN1*-GFP, * p < 0.05, ** p < 0.01, *** p < 0.001, **** p < 10^-4^. Detailed statistical results in **Supplementary Table 1**. Rpm, rotation per minute, a.u., arbitrary units.

We then carried out a bilateral stereotactic injection in the DS of R6/2 mice of AAVs expressing PYK2 and GFP (R6/2-*Camk2a*-PYK2 group) or GFP alone (R6/2-*Camk2a*-GFP group). As control, we injected the *Camk2a*-GFP AAV in wild type mice (WT-*Camk2a*-GFP group). We measured the body weight in the three groups of mice and found a progressive decrease in the R6/2 mice compared to WT, independently of the injected AAV (**Figure 4B**, **Supplementary Table 1**). When we examined hind limb clasping, we observed this response in both R6/2-*Camk2a*-GFP and R6/2-*Camk2a*-PYK2 mouse groups, in contrast to WT-Camk2a-GFP mice, which did not display this response (**Figure 4C**). We then measured the horizontal locomotion of the three groups of mice in a circular corridor maze (**Figure 4D-E**). The general locomotor activity was decreased at 9 weeks in R6/2 mice groups (significant in R6/2 GFP, **Figure 4D**, **Supplementary Table 1**). Rearing activity was significantly decreased at 9 weeks in R6/2-PYK2 mice (**Figure 4E**, **Supplementary Table 1**). We then assessed the potential effects of PYK2 transduction on motor coordination, using the accelerating rotarod (**Figure 4F-G**). The latency to fall (**Figure 4F**, **Supplementary Table 1**) and the speed at fall (**Figure 4G**, **Supplementary Table 1**) were dramatically decreased in R6/2 mice as compared to WT, in both R6/2-*Camk2a*-GFP and R6/2-*Camk2a*-PYK2 mouse groups. Thus, our results showed that PYK2 expression in the DS of R6/2 mice with a *Camk2a* promoter had no effect on their phenotype.

### 3.5. Partial effects of AAV-mediated PYK2 striatal expression under *SYN1* promoter in R6/2 mice

Because of the lack of effect of PYK2 viral expression under the control of *Camk2a* promoter, we tried a different AAV, in which expression was controlled by the human *SYN1* promoter. The procedure was the same as above, with a bilateral stereotactic injection of viral vector into the DS of WT and R6/2 mice. We first compared the effects of the injection of AAV-*SYN1*-GFP in one DS and AAV-*SYN1*-PYK2 in the contralateral DS in WT mice (**Figure 5A**). PYK2 immunofluorescence appeared increased on the AAV-*SYN1*-PYK2-injected side (**Figure 5A**).

**Figure 5:**
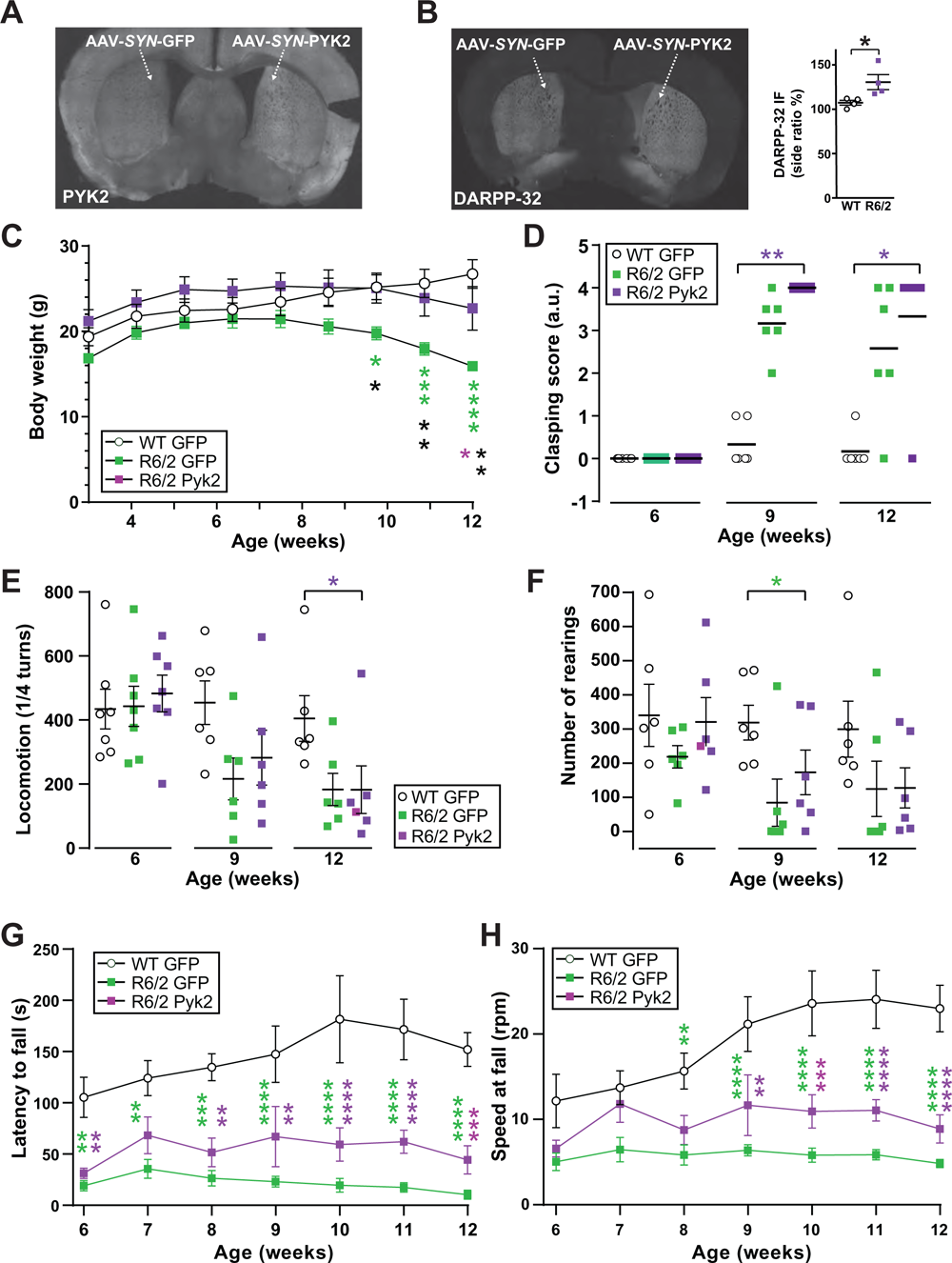
Effects of AAV-*SYN1*-PYK2 transduction in the DS of R6/2 mice. **A**) AAV-*SYN1*-PYK2, expressing PYK2 and GFP, was stereotactically injected in one striatum of a WT mouse and AAV-*SYN1*-GFP, expressing GFP only, in the contralateral striatum. Coronal brain sections were immunolabeled for PYK2. **B**) Four-week-old WT and R6/2 mice were injected as in **A** and DARPP-32 detected by immunofluorescence in coronal brain sections. Left panel, sample picture of a WT mouse. Right panel, immunofluorescence was compared in the right and left striatum of WT and R6/2 mice and the ratio quantified (DARPP-32 IF AAV-*SYN1*-PYK2-injected side / AAV-*SYN1*-GFP-injected side). Mann-Whitney test, p = 0.029 (n = 4). **C**) R6/2 mice received a bilateral injection of either AAV-*SYN1*-PYK2 or AAV-*SYN1*-GFP in the DS and WT mice received a bilateral injection of AAV-*SYN1*-GFP. Their body weight was measured every week (6 mice per group). Two-way repeated measures ANOVA, interaction, F_(16,120)_ = 9.09, p < 10^-4^, age effect, F_(8,120)_ = 18.07, p < 10^-4^, mouse/AAV group effect, F_(2,15)_ = 3.64, p = 0.051. Asterisks indicate significant differences with Holm-Sidak’s multiple comparisons test. **D**) Clasping responses were scored in the 3 groups at the indicated ages (6 mice per group). Kruskal-Wallis test, p < 10^-4^. Horizontal bars are means. **E-F**) The motor behavior of the 3 groups of mice (6 mice per group) was studied in a circular maze at the indicated ages. Locomotion was measured as the number of ¼ turns (**E**) and the number of rearings counted (**F**). Kruskal-Wallis test, (**E**), 6 w, ns, 9 w, ns, 12 w, p = 0.033, (**F**), 6 w, ns, 9 w, p = 0.034, 12 w, ns. **G-H**) The motor coordination was evaluated every week from 6 to 12 weeks in a rotarod test by measuring the latency to fall (**G**) and the speed at fall (**H**). Two-way repeated measures ANOVA (6 mice per group at every age), (**G**) interaction, F_(12,90)_ = 1.94, p = 0.04, age effect, F_(6,90)_ = 2.93, p = 0.01, mouse/AAV group, F_(2,15)_ = 19.8, p < 10^-4^, (**H**) interaction, F_(12,90)_ = 4.74, p < 10^-4^, age, F_(6,90)_ = 7.84, p < 10^-4^, mouse group, F_(2,15)_ = 14.56, p = 0.0003. Asterisks indicate significant post hoc Holm-Sidak’s multiple comparisons test. (**C**, **E-H**) Means ± SEM are indicated. (**D-H**) Post hoc Dunn’s multiple comparisons test. Significant differences are indicated by asterisks, green R6/2 AAV-*SYN1*-GFP vs WT AAV-*SYN1*-GFP, violet, R6/2 AAV-*SYN1*-PYK2 vs WT AAV-*SYN1*-GFP, black, R6/2 AAV-*SYN1*-PYK2 vs R6/2 AAV-*SYN1*-GFP, * p < 0.05, ** p < 0.01, *** p < 0.001, **** p < 10^-4^. Detailed statistical results in **Supplementary Table 1**. Rpm, rotation per minute, a.u., arbitrary units.

We looked for possible effects of AAV-*SYN1*-PYK2 injections in both WT and R6/2 mice on immunofluorescence of DARPP-32 (**Figure 5B**), a protein highly enriched in SPNs. To allow a more accurate evaluation of immunofluorescence, we compared one DS injected AAV-*SYN1*-PYK2 to the other side injected with AAV-*SYN1*-GFP, in the same brain sections in animals of the same genotype. The DARPP-32 immunoreactivity ratio of AAV-*SYN*1-PYK2-injected side vs. AAV-*SYN1*-GFP-injected side was increased in R6/2 mice compared to WT mice (**Figure 5B**). This result suggested a positive effect of PYK2 viral expression on SPNs in R6/2 mice.

We then studied the effects of bilateral striatal injection of AAV-PYK2 on the R6/2 mice phenotype by generating three groups of mice, R6/2 mice injected with *SYN1*-GFP or *SYN1*-PYK2 AAV, and WT mice injected with *SYN1*-GFP AAV as control. We monitored the body weight loss and found that at 10, 11 and 12 weeks of age it was significantly less pronounced in R6/2-*SYN1*-GFP mice than in WT-*SYN1*-GFP mice (**Figure 5C**, **Supplementary Table 1**). This result indicated a beneficial effect of PYK2 transduction on body weight alteration in R6/2 mice. In contrast the clasping score was increased in R6/2 mice without improvement in R6/2-*SYN1*-PYK2 mice (**Figure 5D**, **Supplementary Table 1**). We did not observe any consistent difference in the locomotion and rearing behavior in the circular corridor maze (**Figure 5E-F**, **Supplementary Table 1**), possibly due to a large variability between animals in this test. The performance of R6/2 mice in the rotarod test evaluated by the latency to fall (**Figure 5G**) and speed at fall (**Figure 5H**) was dramatically altered in R6/2 groups compared to WT mice. This alteration appeared to be slightly delayed and consistently less profound in R6/2-SYN1-PYK2 mice than in R6/2-SYN1-GFP mice, without reaching statistical significance between these two groups at any time point (**Figure 5G-H**, **Supplementary Table 1**). Overall our results with the *SYN1*-PYK2 AAV revealed slight positive effects on the weight loss and possibly on motor coordination of R6/2 mice.

## 4. Discussion

### 4.1. PYK2 protein levels are decreased in the striatum of R6/2 mice and HD patients, in part through decreased mRNA

PYK2 is a non-receptor tyrosine kinase highly expressed in the forebrain principal neurons (de Pins et al., 2021). We show that its mRNA levels are higher in the hippocampus than in the cerebral cortex, and intermediate in the striatum, in agreement with previous *in situ* hybridization and immunoblotting data (Menegon et al., 1999). Recent findings showing that PYK2 levels were low in the hippocampus of R6/1 HD mouse model and human HD patients suggested that PYK2 may participate in the pathophysiology of HD in the hippocampus (Giralt et al., 2017). Our results provide the first information about the possible role of PYK2 in striatal HD-like phenotype. As in hippocampus, the striatal levels of PYK2 protein were decreased in R6/2 mice and HD patients. Here we provide evidence in the mouse HD models that the protein deficit is likely to be the consequence, at least in part, of decreased mRNA levels in both hippocampus and striatum. This observation is in agreement with the role of huntingtin in regulating transcription (Saudou and Humbert, 2016). Accordingly alteration of many transcripts were reported in the striatum of HD patients, including PYK2 (*PTK2B* gene) (Hodges et al., 2006). Additional mechanisms, not explored in our study, can potentially contribute to the decrease in PYK2 protein. Cell death is unlikely to be involved since it is very low in the striatum of R6/2 mice in which the resistance of SPNs to malonate-induced cell death is even increased (Hansson et al., 2001). However, the Ca^2+^-activated protease calpain, which cleaves PYK2 (Marzia et al., 2006), is known to be activated in HD cells (Incebacak Eltemur et al., 2022; Paoletti et al., 2008) and could play a role in decreasing PYK2 protein in addition to decreased mRNA.

### 4.2. The absence of PYK2 protein in KO mice does not mimic the HD striatal phenotype

The hippocampal phenotype of PYK2-KO mice was partly similar to that of R6/1 mice, an observation which led to the hypothesis of a contribution of PYK2 deficit in the HD-like hippocampal phenotype (Giralt et al., 2017). Therefore, we explored whether the striatal decrease in PYK2 levels could contribute to the striatal-related HD-like symptoms in mice. We did not observe such phenotype in the complete absence of PYK2 in young KO mice. Accordingly, we previously reported that 3-6 month-old PYK2-KO mice did not display any deficit in the accelerating rotarod task (de Pins et al., 2020). Yet in response to a pharmacological challenge by cocaine or a dopamine D1-receptor agonist, the absence of PY2 decreased the acute locomotor response (de Pins et al., 2020), indicating a contribution of PYK2 in the function of SPNs only revealed in specific conditions. Our results rule out the possibility that PYK2 deficit by itself could be a major player in HD-like striatal manifestations in R6/2 mice, but leave open the possibility that it play a role in modulating striatal neurons responses when they are challenged by pharmacological agents or pathological conditions.

### 4.3. Viral expression of PYK2 in the striatum of R6/2 mice has slight positive effects

Because we hypothesized that PYK2 deficit in striatal neurons could contribute to the HD phenotype, not as a major primary mechanism, but only in conjunction with the other molecular alterations, we investigated whether viral expression of PYK2 in the DS could improve the symptomatology. We had previously shown that hippocampal viral expression of PYK2 rescued some deficits including Y-maze spontaneous alternation and novel object location tests as well as dendritic spine markers, whereas other parameters such as synaptic plasticity were not improved (Giralt et al., 2017). In the current study, we used two different AAV vectors, AAV1-*Camk2a*-eGFP-2A-mPTK2B (AAV-*Camk2a*-PYK2) and AAV9-*hSYN1*-eGFP-2A-mPTKB2B (AAV-SYN1-PYK2). None of them resulted in a massive overexpression but both increased PYK2 levels in control experiments. There was no effect of AAV-*Camk2a*-PYK2 on weight loss or motor phenotype. In contrast we observed a slight improvement of several parameters following bilateral injection of AAV-*SYN1*-PYK2 compared to AAV-*SYN1*-GFP. We do not know the reason for the apparent differences between the phenotypic effects of the two viral vectors we used. Although expression levels appeared similar, functional differences might be related to the different promoters (mouse *Camk2a* vs. human *SYN1*) or AAV serotypes (AAV1 vs AAV9). In support of the latter possibility, a systematic comparison of different brain regions indicated a better neuronal transduction by AAV9 than AAV1 in the striatum (Kofoed et al., 2022). The effect of AAV-*SYN1*-PYK2 was significant for weight loss and a positive trend was visible on the rotarod performance. In addition, an increase in DARPP-32 immunofluorescence was observed in the virally-transduced area indicating a possible positive effect on the biology of SPNs. The effect on weight loss was the most striking, although it was not expected because the weight alteration is usually ascribed to extrastriatal alterations (Bozzi and Sciandra, 2020; Cheong et al., 2019). Further experiments would be needed to test whether it involves an effect on dopamine D1 or D2 receptor-expressing SPNs, which both have been associated with metabolic alterations including changes in body weight in rodents (Johnson and Kenny, 2010; Kim et al., 2014). Together these results suggest a minor but real contribution of PYK2 deficit in the striatum in the phenotype of R6/2 mice.

### 4.4. PYK2 multiple roles in various neurodegenerative and psychiatric conditions

PYK2 emerges as a multifaceted regulator of numerous pathophysiological processes in the brain (de Pins et al., 2021). The current results combined with a number of other studies over the last years allow to position better PYK2 contribution in neuropsychiatric disorders. In HD, our results indicate that PYK2 may contribute to the hippocampal phenotype but plays a lesser role in the striatal phenotype. In Alzheimer’s disease (AD), GWAS meta-analyses showed that the locus of PYK2 gene, *PTK2B*, is associated with an increased risk for late onset AD (Lambert et al., 2013), an association confirmed in several other studies (Beecham et al., 2014; Jiao et al., 2015; Lawingco et al., 2021; Nettiksimmons et al., 2016; Schwartzentruber et al., 2021; Wang et al., 2015). In AD, PYK2 appears to have dual roles in animal models, contributing to the amyloid cascade leading to neurodegeneration (Salazar et al., 2019), whereas it also has a protective role in some mouse models (Brody et al., 2022; Giralt et al., 2018; Kilinc et al., 2020). PYK2 possible role in psychiatric conditions has also been documented, mostly in animal models. Deletion of PYK2 in the amygdala improves some components of depressive-like symptoms induced by a chronic unpredictable mild stress (Montalban et al., 2019), whereas enhancing PYK2 expression in lateral septum neurons can have antidepressant effects (Sheehan et al., 2003). In the context of addiction, PYK2 is induced by chronic cocaine in the striatum and frontal cortex (Freeman et al., 2002; Freeman et al., 2001). In nucleus accumbens neurons expressing D1 receptors PYK2 is necessary for acute locomotor effects of the cocaine but not for its plasticity-related manifestations, locomotor sensitization and conditioned place preference (de Pins et al., 2020), whereas in orbitofrontal cortex PYK2 overexpression prevents the spine loss and behavioral alterations induced by the drug (Whyte et al., 2021). Thus, depending on the brain structures, pathologies, and animal models, PYK2 can play positive or negative roles in neuropsychiatric conditions. In addition, PYK2 is known to be an important contributor to the pathology of various cancers including breast cancer (Gil-Henn et al., 2023) and search for pharmacological inhibitors of selective PYK2 is an active area. Therefore, further study of PYK2 in physiology and pathological conditions is important to balance the potential benefits and risks of its enhancement or inhibition in any of them.

### 4.5. Limitations of the study, therapeutic implications, and conclusions

Our study has some limitations. First, the R6/2 mouse model we used displays a severe phenotype, which starts at an early age. This may have limited potential beneficial effects of PYK2 viral expression as compared to the milder model, R6/1 used in the hippocampal study (Giralt et al., 2017) or to knock in models such as zQ175 (Kacher et al., 2019; Menalled et al., 2012), in which it would be interesting to study the role of striatal PYK2. A second factor was the moderate expression of PYK2 induced by the two viral vectors we used. None of them fully restored PYK2 levels, and this may also have been a limiting factor. Another limitation in the study of PYK2 KO on striatal function is that it was done in adult mice younger than 6 months to compare it with our observations in R6/2 mice and we did not explore whether deficits could occur at an older age.

### 4.6. Conclusions and therapeutic implications

Our study contributes to the evaluation of the multifaceted roles of PYK2 in mouse models of neurological and psychiatric diseases. It reveals a PYK2 deficit in the striatum of HD patients and R6/2 mouse model. It also provides evidence that this defect results at least in part from decreased mRNA levels. The genetic deletion of PYK2 did not result in a motor phenotype by itself. Viral expression of PYK2 in the striatum only led to a slight improvement of the R6/2 mice phenotype. These results show that striatal PYK2 is not necessary for motor behavior and is likely to play only a minor role in the striatal pathophysiology of the HD R6/2 mouse model used. Thus, the role of PYK2 in HD mouse models appears more prominent in the hippocampus than in the striatum. This may be due to the higher expression levels of PYK2 in the hippocampus, where its deletion leads to noticeable alterations (Giralt et al., 2017) although less modifications were reported in a different mouse line (Salazar et al., 2019). An indirect support of a possible contribution of striatal PYK2 in HD is provided by the beneficial effects on motor coordination in R6/1 mice of genetic deletion of the striatal-enriched tyrosine phosphatase, STEP (Garcia-Forn et al., 2018), which dephosphorylates and functionally antagonizes PYK2 [see (de Pins et al., 2021) for a review of PYK2 regulation]. Therefore, in addition to the difficulty to increase PYK2 function, its direct manipulation in the striatum of HD patients does not appear likely to bring sufficient therapeutic benefit. In contrast, an intervention at the level of STEP, which has other relevant targets in addition to PYK2, might be more promising.

## CRediT authorship contribution statement

Omar Al-Massadi: Conceptualization, Formal analysis, Investigation, Methodology, Visualization, Writing-original draft. Benoit de Pins: Formal analysis, Investigation, Methodology. Sophie Longueville: Formal analysis, Investigation, Methodology. Albert Giralt: Investigation, Methodology, Resources. Theano Irinopoulou: Methodology, Formal analysis, Visualization. Mythili Savariradjane: Investigation, Methodology, Visualization. Enejda Subashi: Investigation, Methodology. Silvia Ginés: Resources. Jocelyne Caboche: Resources, Supervision. Sandrine Betuing: Conceptualization, Methodology, Resources, Supervision. Jean-Antoine Girault: Conceptualization, Data curation, Formal analysis, Funding acquisition, Project administration, Resources, Supervision, Validation, Writing - review & editing.

## Data availability

Data will be made available on request.

### Abbreviations

AAV: adeno-associated virus
AD: Alzheimer’s disease
ANOVA: analysis of variance
a.u.: arbitrary unit
CAG: cytosine-adenine-guanine
DARPP-32: dopamine-and cAMP-regulated phosphoprotein with an apparent molecular weight of 32 kDa
DS: dorsal striatum
GWAS: genome-wide association study
HD: Huntington’s disease
KO: knockout
ns: not significant
PBS: phosphate-buffered saline
PBS-T: PBS with Triton X-100
PMI: post mortem interval
*PTK2B*: protein tyrosine kinase 2B, gene coding for PYK2
PYK2: proline-rich tyrosine kinase 2
rpm: rotation per minute
RT-qPCR: real time quantitative polymerase chain reaction
SPN: spiny projection neuron
STEP: striatal-enriched tyrosine phosphatase

## Supporting information

Results of statistical analyses

## Acknowledgments

We thank the *Mouse breeding and phenotyping facility* of the *Institut du Fer à Moulin* (IFM) for animal care and breeding and the IFM *Cell and Tissue Imaging facility*, where image acquisitions were performed.

